# Characterization of the Pace-and-Drive Capacity of the Human Sinoatrial Node: a 3D in silico Study

**DOI:** 10.1101/2022.06.03.494644

**Authors:** Antoine Amsaleg, Jorge Sánchez, Ralf Mikut, Axel Loewe

**Affiliations:** Institute of Biomedical Engineering, Karlsruhe Institute of Technology (KIT), Karlsruhe, Germany; Institute for Automation and Applied Informatics, Karlsruhe Institute of Technology (KIT), Karlsruhe, Germany

## Abstract

The sinoatrial node (SAN) is a complex structure that spontaneously depolarizes rhythmically (“pacing”) and excites the surrounding non-automatic cardiac cells (“drive”) to initiate each heart beat. However, the mechanisms by which the SAN cells can activate the large and hyperpolarized surrounding cardiac tissue are incompletely understood. Experimental studies demonstrated the presence of an insulating border that separates the SAN from the hyperpolarizing influence of the surrounding myocardium, except at a discrete number of sinoatrial exit pathways (SEP). We propose a highly detailed 3D model of the human SAN, including 3D SEPs to study the requirements for successful electrical activation of the primary pacemaking structure of the human heart. A total of 788 simulations investigate the ability of the SAN to pace and drive with different heterogeneous characteristics of the nodal tissue (gradient and mosaic models) and myocyte orientation. A sigmoidal distribution of the tissue conductivity combined with a mosaic model of SAN and atrial cells in the SEP was able to drive the right atrium (RA). Additionally, we investigated the influence of the SEPs by varying their number, length and width. SEPs created a transition zone of transmembrane voltage (TMV) and ionic currents to enable successful pace and drive. Unsuccessful simulations showed a hyperpolarized TMV (−66 mV), which blocked the L-type channels and attenuated the sodium-calcium exchanger. The fiber direction influenced the SEPs that preferentially activated the crista terminalis (CT). The location of the leading pacemaker site (LPS) shifted towards the SEP-free areas. LPSs were located closer to the SEP-free areas (3.46±1.42 mm), where the hyperpolarizing influence of the CT was reduced, compared to a larger distance from the LPS to the areas where SEPs were located (7.17±0.98 mm). This study identified the geometrical and electrophysiological aspects of the 3D SAN-SEP-CT structure required for successful pace-and-drive in silico.

**SIGNIFICANCE:** The human sinoatrial node (SAN) is the intrinsic natural pacemaker of the heart. Despite its remarkable robustness to failure, the electrophysiological properties, and mechanisms by which the SAN overcomes the source-sink mismatch towards the hyperpolarized surrounding cardiac tissue remains a mystery. The SAN is electrically isolated from the hyperpolarized cardiac tissue, except at a discrete number of sinoatrial exit pathways (SEP). Using in silico experiments, we explore the influence of the fiber orientation, the SEPs’ number, geometry and location on the activation of the SAN and the surrounding atrial tissue. We provide the mechanisms in a first 3D model of the human SAN-SEP structure that can successfully drive the working myocardium.

## INTRODUCTION

The sinoatrial node (SAN) is the primary natural pacemaker of the heart, responsible for its electrical activation (1–4). The SAN is located across the entire superior vena cava region between the crista terminalis (CT) and the interatrial septum. The SAN is connected to the CT as part of the non-automatic working myocardium of the right atrium (RA) (1, 5–12). The CT is a muscular bundle involved in the rapid electrical propagation of the excitation that initiates in the SAN and spreads across the RA. The conduction velocity (CV) in the CT is high (between 100 cm/s to 140 cm/s) compared to other regions of the RA (approximately 70 cm/s) and the SAN (between 5 cm/s and 20 cm/s) (2, 13–15) to allow for a rapid spread of the excitation across the whole RA. The big difference in conductivity between CT and SAN needs a delicate transition to protect the SAN from the hyperpolarizing effect of the CT that acts as a sink for the current generated by the SAN.

The SAN is a complex structure of specialized cardiac tissues different from the surrounding atrial myocardium (1, 16–18), which spontaneously and rhythmically triggers an electrical impulse (pace). The mechanism responsible for the SAN “pace” behavior has been identified as a delicate balance of ionic currents leading to the spontaneous depolarization of the SAN cells known as the “coupled clock mechanism” (17, 19, 20). The electrical depolarization wave, originated in the SAN, then propagates to the surrounding cells (drive) and leads to the contraction of the heart. However, the SAN can fail to pace because of the hyperpolarizing influence of the CT/RA myocytes, which suppresses automaticity in the SAN cells. When pacing, it can furthermore fail to drive the CT when the depolarization wave is blocked at the exit of the SAN (exit block). The mechanisms by which the small SAN is able to pace and drive the large and hyperpolarized surrounding cardiac tissue, without being suppressed, are far from being understood (3, 6, 8, 21, 22).

Previous studies showed that the SAN is electrically isolated from the surrounding myocardium, except at a discrete number of conducting channels known as the sinoatrial exit pathways (SEP) (2, 3, 8, 23–25). This SAN-SEP structure, combined with heterogeneous properties, make the SAN robust to dysfunction (8, 23). Indeed, the SAN requires anatomical (fibroblasts, collagen, adipose tissue and blood vessels) (3, 8, 10, 18, 24, 26) and functional heterogeneity (sparsity of connexins) (7, 27) to protect it from the hyperpolarizing influence of the CT (8) in order to maintain its physiological function as the leading pacemaker. Different studies have proposed two models (gradient or mosaic) of the heterogeneous SAN and its connection to the myocardium (1, 6–8, 18, 19, 24, 27–30). The gradient model describes a change of conductivity from the SAN center to its periphery to represent the sparsity of connexins in the SAN and achieve a gradual change of CV from the SAN center via the SEP to the CT. The mosaic model was proposed to explore the effect of cellular heterogeneity (mix of different densities of SAN cells, fibroblasts and atrial myocytes) present in the SAN (6, 27, 29, 31). However, both have been explored separately, which likely oversimplifies the complex structure of the SAN and the cellular interaction. Additionally, the cell’s orientation along the SAN structure changes progressively from the inner SAN towards the CT (6, 7).

The SAN cellular characteristics have been extensively studied (16, 17, 19). However, the characterization of the cardiac myocytes in the CT, that interact directly with the nodal tissue, has not been fully explored. Several computational studies have proposed modifications of different ionic channels maximum conductance to reach a value close to the experimentally reported repolarization time (15, 29, 32, 33). Moreover, experimental data from CT myocytes reported that their upstroke velocity is faster than in other regions of the RA (34).

In silico experiments help better understand the effect of the cellular heterogeneity present in the SAN-SEP structure and its electrophysiology at the tissue level. Furthermore, they allow exploring a range of parameters providing insight into the mechanisms of the electrical depolarization, initiated in the SAN and propagated to the surrounding cardiac tissue. However, the current models of the SAN did not consider the 3D structure of the SEPs (24, 25, 35) and studied the gradient and mosaic models separately (7, 8, 29, 30).

Using in silico experiments, we explore the electrophysiological and structural characteristics of the SAN-SEP-CT complex required for successful pace and drive. We aim to quantify the three-dimensional composition of the SAN-SEP at the cellular and tissue level and the effect on the depolarization of the cardiac tissue. Additionally, we aim to explore the effect of the geometrical variability of the SEPs.

## MATERIALS AND METHODS

### Electrophysiology model

To represent the electrophysiology of the human right atrial myocardium, we used the mathematical model proposed by Courtemanche et al. (36). To represent the electrophysiological heterogeneity of the RA, we modified the Courtemanche et al. model as proposed by Wilhelms et al. (29) to reproduce the repolarization time of the cardiac myocytes in the CT. To reproduce the prolongation of the action potential duration (APD) observed in the CT, two ionic channel maximum conductance were decreased: the rapid delayed rectifier potassium channel (G_Kr_) and the transient outward potassium channel (G_to_) were decreased by 50%. Additionally, we increased the maximum conductance of the sodium (G_Na_) by 100% to reproduce the action potential (AP) upstroke velocity reported by Avanzino et al. (34). The electrophysiology of the human SAN was represented by the mathematical model described by Loewe et al. (37, 38).

### Tissue model

The SAN was modeled as an ellipsoid, 20 mm long, 4 mm wide and 2.6 mm thick (10, 18, 39) (Figure 1B and C). As the SAN is surrounded by fibrotic and adipose tissue, our model represents this by a cavity that isolates the SAN from the cardiac tissue. The SEPs were modeled by cylinders with a diameter of 0.6 mm and a length of 1.5 mm as experimental data on SEP geometry is scarce. The SEPs cross the insulating border but also extend in the cardiac tissue (3), which is represented in the model by semi-spherical SEP extensions into the CT of 1 mm radius (Figure 1C). The CT was modeled by a 5 mm wide strip, laterally surrounded by two 5 mm strips of common right atrial tissue (Figure 1A) of 0.8 mm thickness (14). The posterior atrial wall curvature was modeled by bending the CT and the surrounding atrial tissue by 2 mm. Figure 1 depicts the complete fully parameterized model, with colors associated to main regions (Figure 1D), developed in Gmsh version 4.8.4 (40). The tetrahedral mesh elements had an average length of 150 *μ*m.

**Figure 1:**
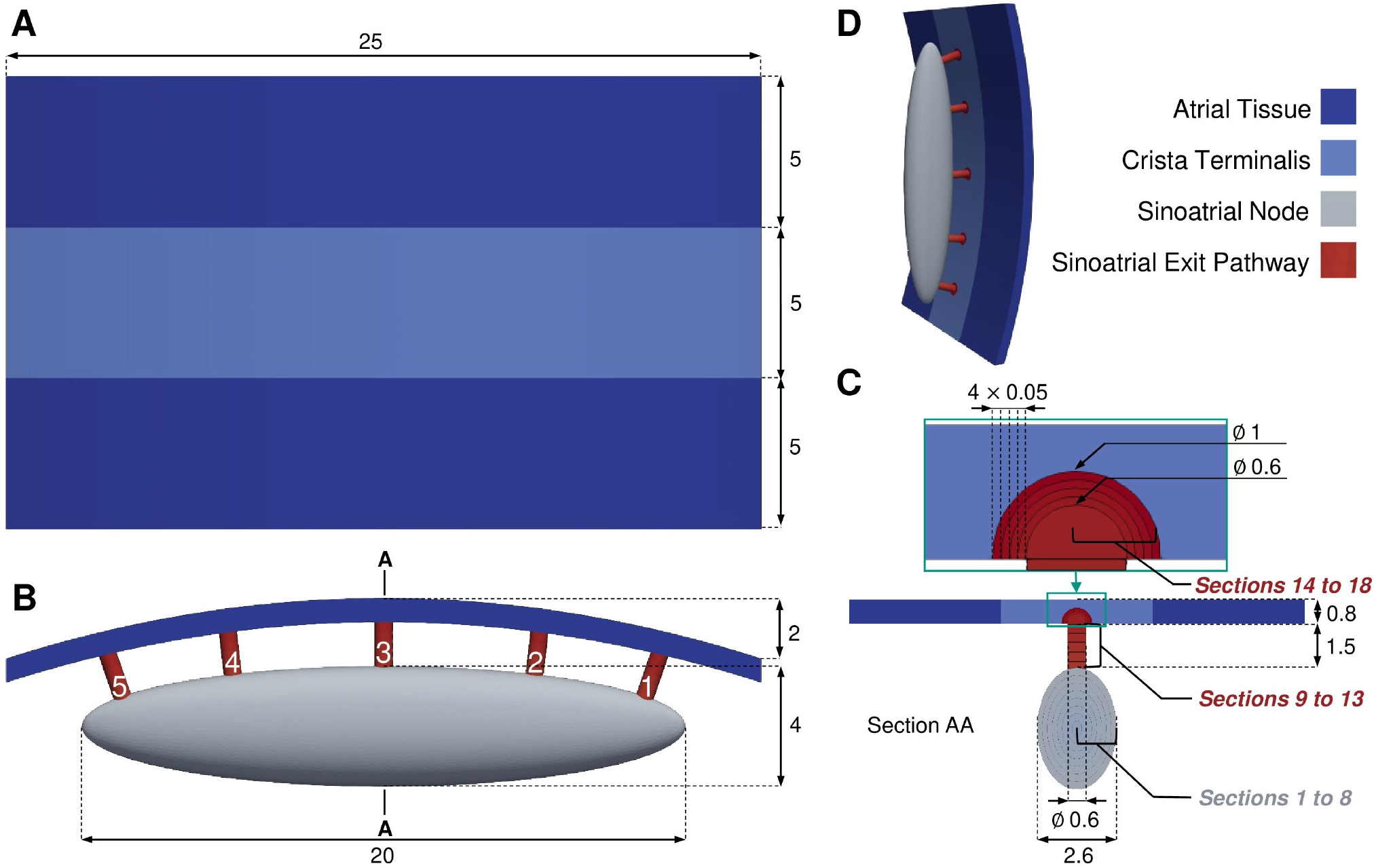
A) Top, B) front, C) side and D) isometric view of the sinoatrial node (SAN) and sinoatrial exit pathways (SEP) model connected to the crista terminalis (CT) and bulk atrial tissue. The SEPs are numbered from 1 to 5 from right to left. The SAN and the SEPs were divided into 8 and 10 sections, respectively (Section AA, panel C). All dimensions are given in millimeters.

To facilitate the study of the electrophysiological properties of the SAN-SEP structure, a reduced model of a single SEP connected to a SAN tissue patch on one side and to a 1 mm × 1 mm CT tissue patch on the other side was built. This simplified model was used to investigate a large number of combinations of cellular and tissue properties to sample the parameter space comprehensively for combinations successfully depolarizing the CT tissue. The size of the CT tissue patch was then increased to 5 mm × 10 mm as the area that a single SEP should activate to make the SAN drive the entire atria. The CT is about 25 mm long (Figure 1A), thus each of the 5 SEPs should activate a fifth of the total CT length (i.e., 5 mm). A safety factor of 2 was applied on top.

### Nodal tissue heterogeneity

The SAN and the SEPs were divided in different sections (Figure 1, Section AA) to implement the electrophysiological changes from the SAN center to its periphery. 8 sections were created in the SAN and 5 in the cylindrical part of the SEP to have a distance between two sections of 0.25 mm to 0.30 mm. In addition, 5 more sections were created at the intersection of the SEP with the CT, as a semi-spherical extension. The distance between the sections was 0.05 mm to adjust the electrophysiological properties at the contact zone with CT more fine grained. In each section, from the SAN center to the SEP extension, a proportion of CT myocytes was defined according to the mosaic model. Moreover, a gradient of conductivity was set up from the central SAN section to the last SEP extension section. These distributions of CT/SAN myocyte density and tissue conductivity were controlled using a sigmoidal distribution function:

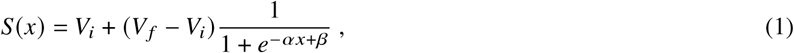

with *x* being the normalized distance from the SAN center to the last SEP extension section, *α* the slope at the inflection point and *β* modulating the abscissa of the inflection point. *V*_*i*_ and *V*_*f*_ are the initial and final values of the distribution, respectively. We set the monodomain tissue conductivity (Table 1) such that the CV of a planar wave in a tissue strand along the fiber direction (41) is 5 cm/s in the SAN center (2, 6, 24), 65 cm/s in the RA tissue and 100 cm/s in the CT (15). The conductivity gradients were set from the SAN center to the CT following Eq. 1. The proportion of CT myocytes ranged from 0% in the first SAN section to 100% in the CT tissue according to Eq. 1.

**Table 1:**
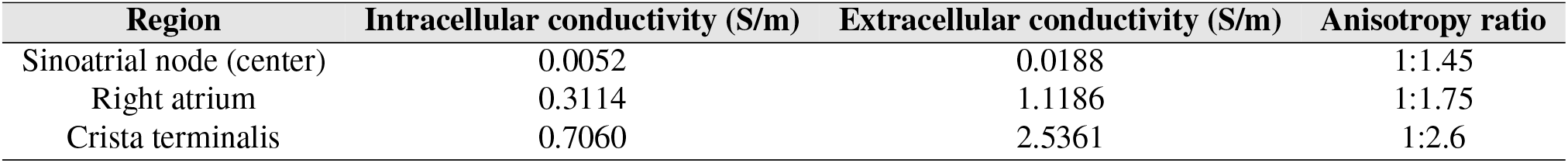
Intracellular and extracellular conductivity for different regions of the atrial tissue along fiber direction and anisotropy.

A full factorial analysis on the four sigmoidal factors of the gradient and mosaic distribution functions was performed in the geometrically reduced model. Five different *α* and *β* values were tested: 0.1, 0.3, 0.5, 0.7, 1.0 and -5, -2, 0, 2, 5 for the gradient and mosaic model, respectively to include a large range of distributions (5^4^ = 625 simulations). The combinations of parameters that successfully depolarized the CT were translated to the complete model.

Furthermore, fiber direction was implemented to represent the gradual orientation change of the cells from the center of the SAN towards the CT (6, 7) (Figure 6Ai). The fiber direction was also symmetrically reversed in a supplementary simulation to prove the effect of the fiber direction on the depolarization of the cardiac tissue (Figure 6Bi).

### Number, length & width of sinoatrial exit pathways

To study the influence of the SEP characteristic on pace and drive behavior, we first removed one SEP at a time. With the fixed locations of the five SEPs, all 26 possibilities were explored with the number of SEPs ranging from 2 to 5 (23): 10 combinations with 2 and 3 SEPs, 5 with 4 SEPs and 1 with 5 SEPs (Figure 2). The local activation time (LAT) was obtained as the time of the maximum upstroke velocity of the last AP triggered by the SAN. The cells that are activated within the first 6 ms, which covers the depolarization phase of the first activated SAN cells, were considered as the leading pacemaker site (LPS). The Euclidean distance from the LPS centroid to two different areas was calculated.

**Figure 2:**
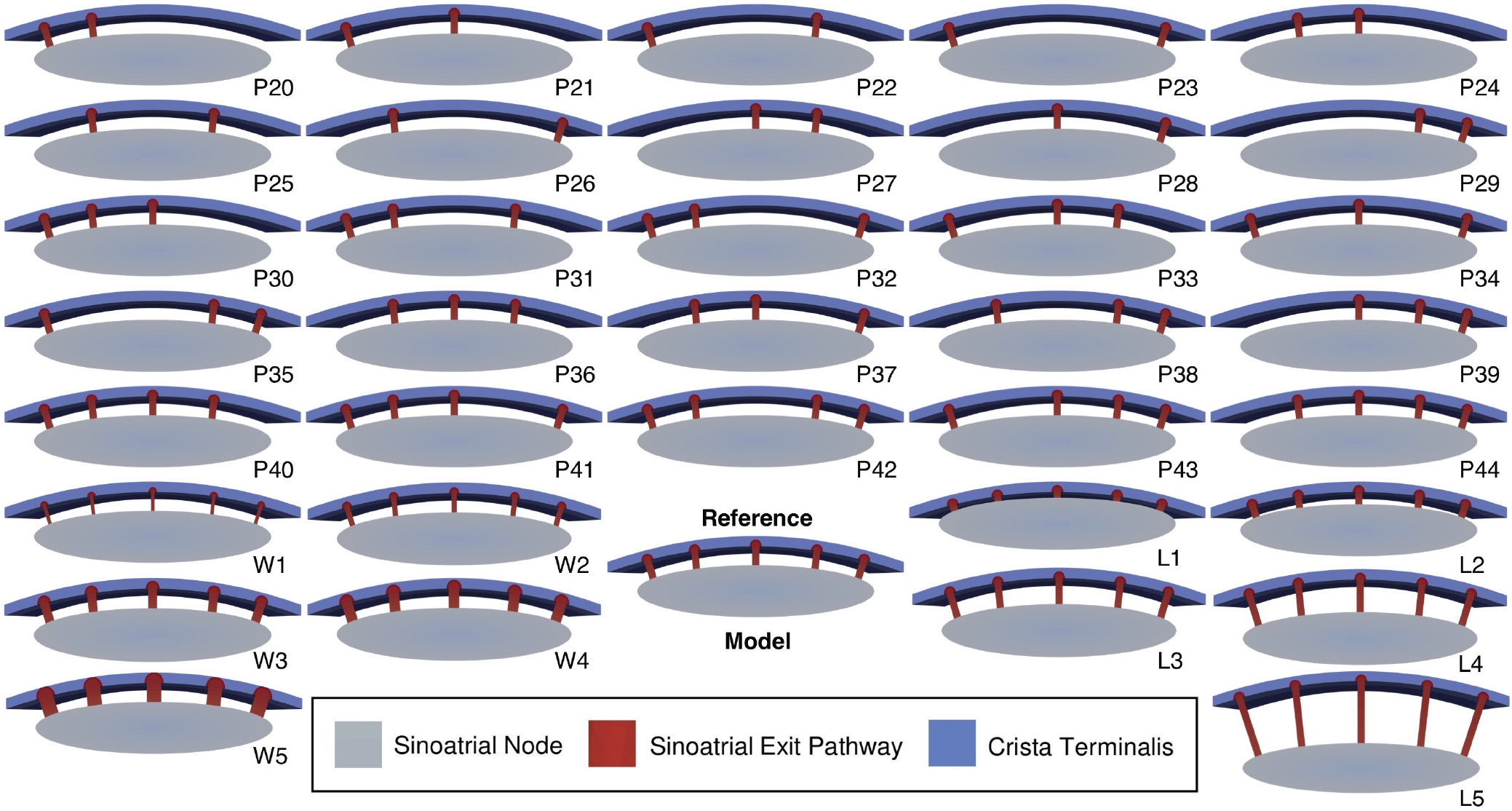
Clipped view of the 36 different sinoatrial exit pathway (SEP) configurations. The reference model comprises 5 SEPs, 1.5 mm long and 0.6 mm wide. P2X, P3X and P4X correspond to the models with 2, 3 and 4 SEPs, respectively. LX and WX correspond to different SEP lengths (0.5, 1, 2, 2.5, 5 mm) and widths (0.3, 0.45, 0.75, 0.9, 1.2 mm), respectively.

We distinguished the SEP areas from the SEP-free areas. SEP areas were defined by one SEP, or groups of consecutive neighbouring SEPs present in the mesh. The SEP-free areas were defined by one removed SEP, or groups of removed consecutive neighbouring SEPs.

For an area with an even number of SEPs, or removed SEPs, the middle of the two central SEPs was calculated. In the odd case, the position of the central SEP was considered. Then, when more than one distance from the LPS centroid to a SEP area or SEP-free area was measured, the mean distance was calculated from the LPS centroid to the previously described location. This aimed to compare the LPS distance to the regions with high CT influence (SEP areas) and to the regions with less CT influence (SEP-free areas). 10 additional simulations investigated the limit dimensions (length, width) of the SEPs and the influence on CT activation. Considering the reference model with SEPs 0.6 mm wide and 1.5 mm long, the SEP dimension variations were 0.5, 1, 2, 2.5, 5 mm for the length and 0.3, 0.45, 0.75, 0.9, 1.2 mm for the width. The 36 different meshes are shown in (Figure 2).

### Numerical simulations

Detailed biophysical monodomain simulations were run using openCARP (42). Simulations ran for 5,000 ms where the first depolarization of the tissue was ignored for the analysis. After reaching a limit cycle, the SAN was able to pace at a constant cycle length (CL). The APs along the depolarization pathway, from the LPS of the SAN to the end of the SEP in the CT were analyzed. The necessary files to run the in silico experiment are available under the Apache License 2.0 (43).

## RESULTS

### Crista terminalis electrophysiology

Different regions of the RA exhibited distinct APD and maximum upstroke velocity (dV/dt_max_) in single cell in silico experiments. The APD of the RA cell was 298.51 ms, 316.63 ms in the CT cell and 304.65 ms in the CT cell with 100% increased sodium channel conductance. Looking at the depolarization phase of the AP, the upstroke for the RA and CT are similar with maximum velocities of 197.79 V/s and 195.17 V/s, respectively. After increasing the sodium channel conductance, dV/dt_max_ was increased to 319.49 V/s.

### Reference model

Among the 625 reduced model simulations, 42 reduced models with a combined gradient and mosaic distribution yielded an activation of the small CT tissue patch (1 mm × 1 mm). However, when the CT dimensions were increased (5 mm × 10 mm) in the second reduced model, only one gradient (*α* = 1, *β* = 2) and mosaic (*α* = 1, *β* = 5) combination (Figure 3A) led to a complete depolarization of the CT with the 100% increased sodium channel conductance. 0% and 50% increase of sodium channel conductance resulted in exit block of the SEPs. Concerning the CT/SAN myocyte distribution, only 201 CT cells were modeled in the transition area compared to 260,366 SAN cells. The increase was observed from the two last sections of the SEP body (sections 12 and 13), before the semi-spherical extension in the CT, with 7% and 15% CT cell share, respectively.

**Figure 3:**
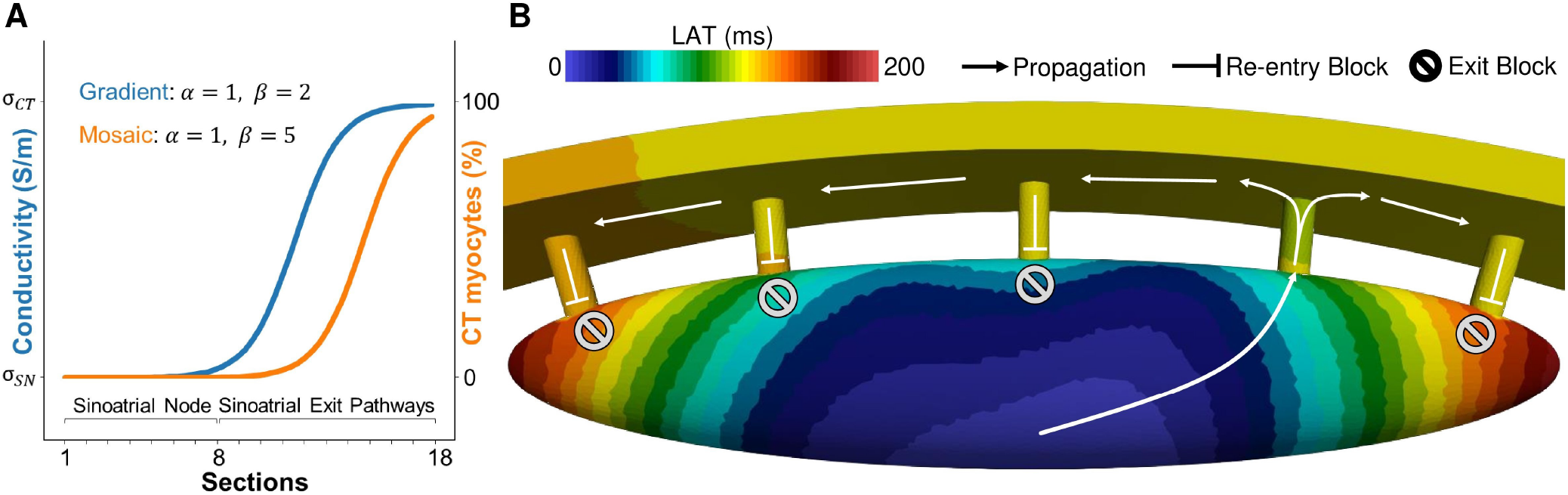
A) Gradient of conductivity and mosaic of crista terminalis (CT) / right atrium (RA) myocyte distributions in the complete sinoatrial node (SAN) and sinoatrial exit pathways (SEP) model yielding successful pace and drive. *α* and *β* are the parameters of the sigmoidal distribution (Eq. 1). The gradient model was implemented distributing the conductivity values from the SAN center to the CT. The CT myocytes were distributed from 0% in the SAN center to near 100% in the CT tissue area. B) Local activation time map of the reference model, showing the activated SEP, the exit block occurring at the other SEPs and the retrograde activation (re-entry) block from the other SEPs to the SAN.

The latter gradient and mosaic distribution was applied to the complete SAN-SEP model in which it also successfully paced and drove the right atrial and CT tissue. The heterogeneous properties of the nodal tissue changed rapidly in the SEP compared to the SAN (Figure 3A). The conductivity increased only in the SEP. The complete SAN (sections 1 to 8) had the same conductivity. As a consequence of the SAN-SEP-CT structure CV was different from its theoretical value, which was tuned for a plane wave in a tissue strand (Figure 4C). Within the LPS area, CV was fast (180 cm/s) due to the cluster of first and simultaneously activated pacemaker cells. The CV then lowered rapidly down to 9 cm/s at the LPS border. Additionally, a small decrease of the CV was observed at the beginning of the transition area, along the path of depolarization, from 5 cm/s down to 1 cm/s. After that, a rapid increase of the CV was observed until the middle of the SEP (section 12) and a second decrease of CV at the SEP insertion (section 14) to then rapidly increased again to the CT CV (124 cm/s).

**Figure 4:**
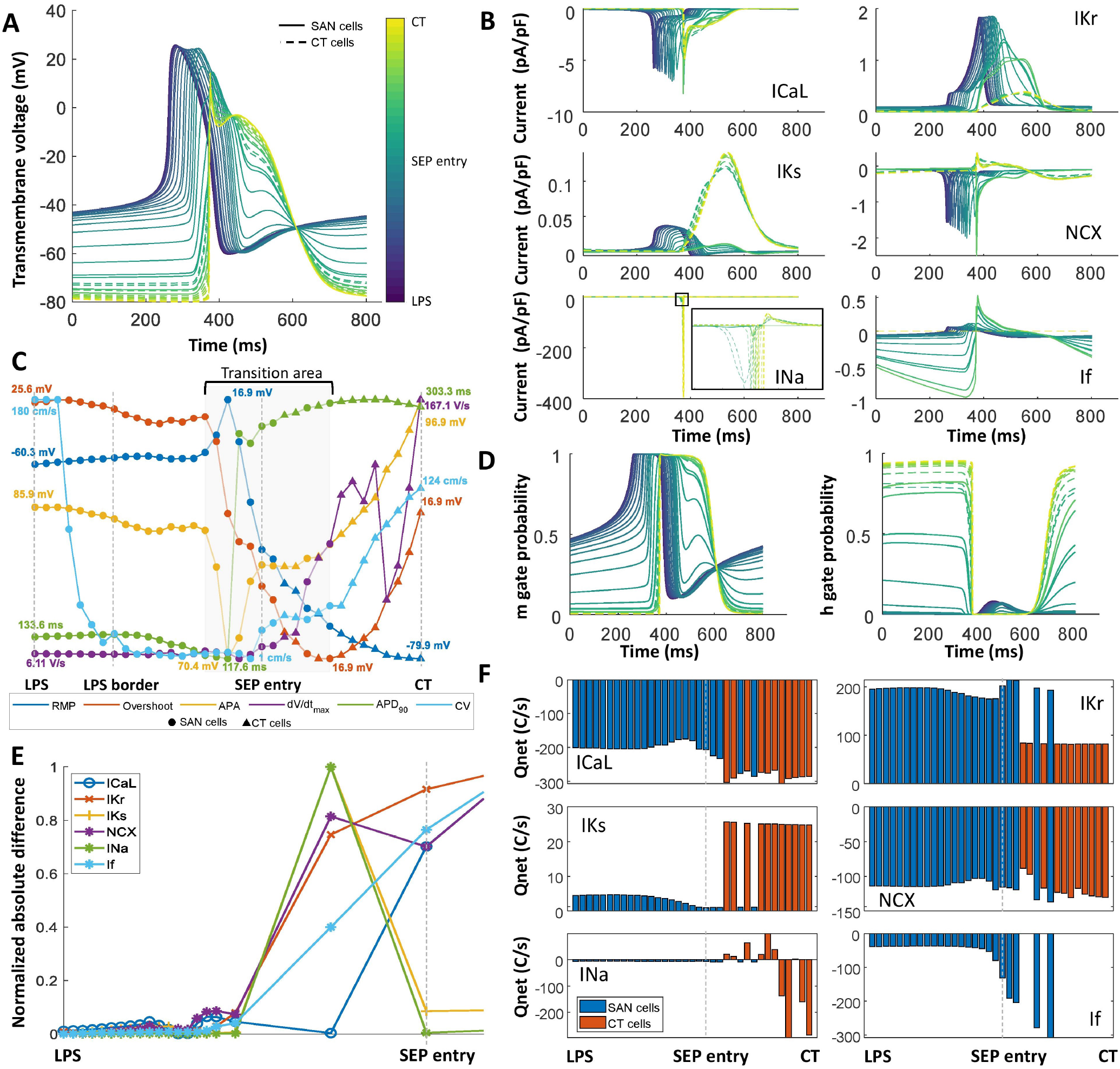
A) Transmembrane voltage evolution from the leading pacemaker site (LPS) to the crista terminalis (CT) across the sinoatrial exit pathway (SEP). B) Current traces for the L-type calcium current (ICaL), rapid delayed potassium rectifier current (IKr), slow delayed potassium rectifier current (IKs), sodium-calcium exchanger (NCX), sodium current (INa) and hyperpolarization-activated current (If). C) Biomarkers measured at different locations along the depolarization path from the LPS to the CT. D) Sodium channel activation (m) and inactivation (h) gate probability. E) Normalized absolute difference between the reference model and model P41. F) Net charge flux of different currents across the cell membrane.

Average LPS AP features were measured for four APs. The mean LPS basic CL was 883±1.73 ms. The maximum diastolic potential in the SAN LPS was -60.30±0.05 mV, the overshoot 25.59±0.04 mV, the AP amplitude 85.90±0.09 mV, the dV/dt_max_ 5.40±0.27 V/s, the APD_90_ of 133.78±0.25 ms and the DDR_100_ was 66.48±0.57 mV/s. In the CT, the resting membrane potential was -80.28±0.25 mV, the 19.46±6.21 mV, the AP amplitude 99.74±6.37 mV, dV/dt_max_ 200.10±2.26 V/s and the APD_90_ was 303.95±1.35 ms. The CV in the CT was 124.62±1.50 cm/s measured between two points in the CT above SEP n^°^3 and SEP n^°^5. The resting membrane potential in the SEP (comprising both CT and SAN cells) ranged between -64.30 mV and -78.99 mV, the overshoot between 8.20 mV and 16.92 mV and the AP amplitude between 72.50 mV and 95.74 mV. The LPS, composed of the pacemaker cells that first activated the SAN had a volume of 6.72 mm^3^, which represented around 6.17% of the total SAN volume. The LAT at the base of the SEP was 87 ms. The maximum LAT of the whole SAN, reached in the ellipsoid extremities, was 174 ms (Figure 3B).

### Pace-and-drive mechanisms

In the reference model, 35 points, starting from the LPS centroid across the SAN and through the activated SEP towards the CT, were traced to look at the evolution of the transmembrane voltage (TMV) and the cellular ionic currents. A change in morphology of the TMV was observed along the depolarization path. In the LPS, the AP of SAN cells did not change compared to the reported values from single cell experiments. Figure 4A depicts different AP from the LPS centroid towards the CT. On the one hand, due to the sink-source effect of the cells distributed in the different SEP sections, CT cells exhibited an AP with a SAN morphology. On the other hand, the SAN cells distributed in the SEP showed a transition of the AP similar to a CT cell. Figure 4C depicts how SAN cells in section 6, 7 and 8 were affected just before the SEP entry (section 9) and followed an evolution to CT biomarker values.

Figure 4B depicts the effect of the sink-source interplay on the ionic currents. The evolution of the L-type calcium current (ICaL), rapid delayed potassium rectifier current (IKr), slow delayed potassium rectifier current (IKs) and sodium-calcium exchanger (NCX) of the SAN cells in the SEP led to a decrease of the net charge during one cycle (Figure 4D), which mainly affected the repolarization phase. Additionally, the CT cells present in the SEP exhibited a more positive resting membrane potential, which led to a partial activation of the fast inactivation (h) gate of the sodium channel. SAN cells in the last sections of the SEP exhibited an increase of the If current due to the prolonged repolarization, which increased the efflux of potassium from the cell. (Figure 4F).

There was a change of sink-source effect during different phases of the cellular depolarization and repolarization. During the AP upstroke in the SAN, the SAN acted as a current source and the CT as a current sink. During the repolarization of the CT cells, the CT acted as a current source to the SAN, which mainly affected the parts of the SEP close to the CT (sections 12 and 13).

The mechanisms underlying non-successful driving of the SAN-SEP were explored in two meshes with 4 SEPs (model P40 and model P41) and with 5 shorter SEPs (model L2). Figure 4E shows that the normalized absolute error between the reference model and model L2 was less than 0.1 before the SEP entry where the SAN was still able to pace in all cases but it was not able to drive the CT. We observed a hyperpolarization of the TMV of cells located at the first SEP section (section 9) due to the sink effect of the CT. The hyperpolarization of the TMV (−66 mV) prevented the ICaL channel from opening, inactivating NCX and not being able to depolarize the SEP cells contributing to the block of conduction in the exit pathways.

### Leading pacemaker site

LPS location was derived as the centroid coordinates of the LPS. Considering all PX models (Figure 5A, B and C) 2,458±844 mesh elements formed the LPS (range: 862 to 4,404 elements). This represented a volume of 5.31±1.77 mm^3^ (range: 1.96 to 9.35 mm^3^), thus around 5% of the whole SAN volume.

**Figure 5:**
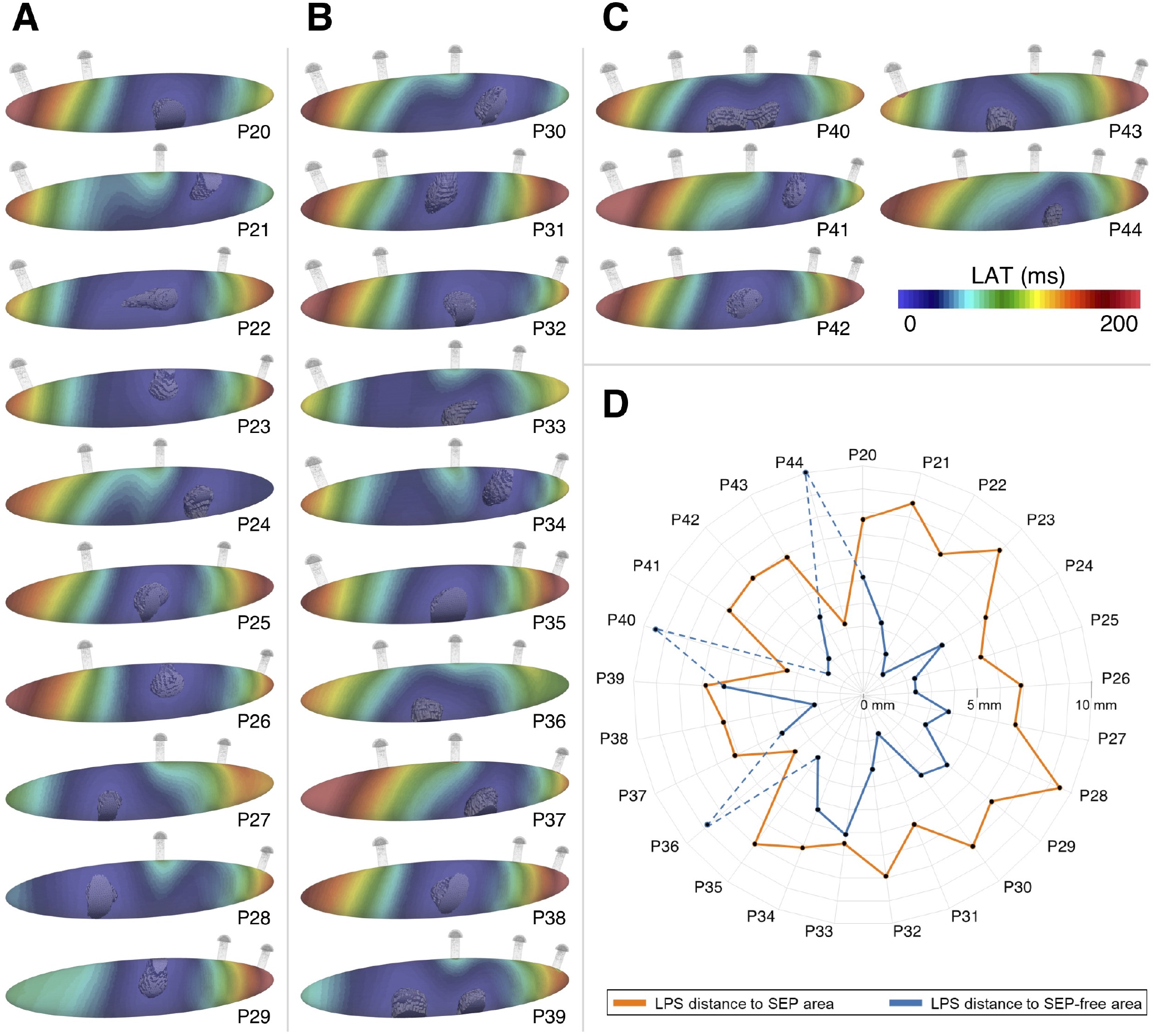
Different combinations of number and location of SEPs with the LATs of the SAN and the corresponding leading pacemaking site (solid gray). Early activated SAN regions are represented in blue and latest activated SAN regions are represented in red. A) Models with 2 SEPs. B) Models with 3 SEPs. C) Models with 4 SEPs. D) LPS distance from the SEP area and SEP-free area. Due to the location of the SEPs in simulations P36, P40 and P44 and the too small volume available at the ellipsoid extremities, the CT influence prevented the LPS to shift.

A shift of the LPS was observed depending on the SEP distribution (Figure 5). The LPS was closer to the SEP-free areas. No LPS was located in the extremities of the ellipsoid, unless when two extreme SEPs (n°1 and n°2 or n°4 and n°5) were removed.

Indeed, if only SEP n°1 or n°5 were removed, this left a smaller SEP-free area for the LPS to shift combined with a still strong influence of the CT because of the presence of SEP n°2 or n°4, respectively. Therefore, the simulations P36, P40 and P44 presented outlier distances from LPS to SEP-free areas. Not considering these three outliers, the LPS was on average located at a distance of 7.17±0.98 mm from the SEP areas and at a distance of 3.46±1.41 mm from the SEP-free areas (Figure 5D).

Full SAN activation from the LPS to the extremities of the ellipsoid took 185±31.59 ms. The depolarization wave reached the first activated SEP n°1 (models P37, P42, P43 and P44) in 174.75±52.44 ms and SEP n°3 (models P28 and P39) in 64.50±2.12 ms.

### Nodal fiber direction

The influence of the fiber direction was evaluated in the reference model. For the reference model (Figure 6Ai), SEP n°2 was activated (Figure 6Aii). When the fiber direction was changed (Figure 6Bi) the symmetrically opposite SEP n°4 was activated (Figure 6Bii). The LAT for the reverse fiber direction was increased by 3.45% to 180 ms with a 60.27% smaller LPS volume of 2.67 mm^3^.

**Figure 6:**
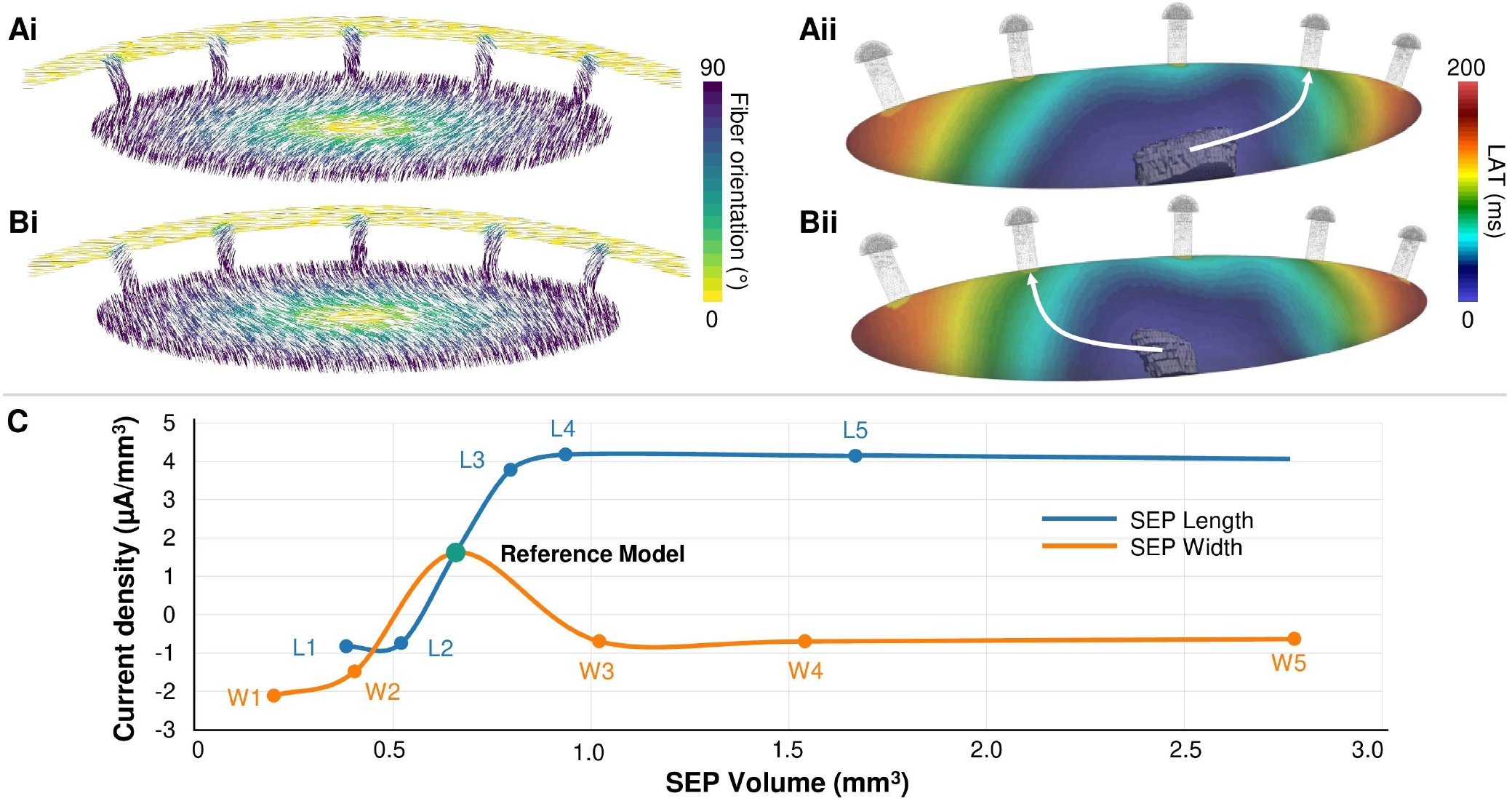
Analysis of the fiber direction and sinoatrial exit pathway (SEP) geometrical dimensions. Ai) Reference model with default fiber direction. Aii) LAT map of the SAN reference model with activation path along SEP n^°^2. Bi) Reference model with reversed fiber direction. Bii) LAT map of the SAN reversed fiber direction model with activation path along SEP n^°^4. C) Evolution of current density as a function of SEP volume.

### Sinoatrial exit pathway dimensions

Starting from the reference model, ten different SEP geometry scenarios were tested. The SEP’s width and length were varied yielding different SEP volumes (Figure 6C). When the width and length of the SEPs was decreased (models W1, W2, L1 and L2) the current generated by the SEP which depolarized the CT was not able to drive the CT. Moreover, when the SEPs width was increased compared to the reference model, the current dropped due to an increase of the contact area of the SEP with the CT. Due to the reduced current across the SEP, the CT tissue was not depolarized.

When the length of the SEPs was increased compared to the reference model, the cardiac tissue still captured the excitation (models L3, L4 and L5). However, for models L3, L4 and L5, the CL was reduced (876 ms, 873 ms and 867 ms, respectively) compared to the reference model.

## DISCUSSION

In this study, we presented a 3D model of the complex SAN-SEP structure, which allows investigating the pace-and-drive capacity of the surrounding myocardium under different conditions. Using in silico experiments, we provide (i) an understanding of the electrophysiological characteristics of the pace-and-drive mechanisms of the human SAN, (ii) insight into the role that SEPs play in the depolarization of the cardiac tissue, (iii) an explanation for the LPS location and role in depolarization.

### Sinoatrial node pace and drive mechanisms

Based on the results of the reference model, we observed the presence of a transition area that drove the depolarization from the SAN LPS to the CT. The transition area started close to the SEP entry, where the cells in the SAN exhibit a change of the net charge across the ICaL and IKs channel. The reduction IKs prolonged the APD (Figure 4C), which also had an impact on the increase of the CL from 883 ms to 887 ms. Verkek et al. observed the effect of the reduction of IKs in patients with heart failure, which were correlated to the change of the CL (44). The observed partial activation of the fast inactivation sodium gate was associated with the driving mechanism that allows the CT to depolarize. SAN dysfunction has been associated with mutations of the sodium channel (45). In the future, the proposed model could be used to investigate the mechanisms behind sodium channel mutations and SAN dysfunction. The failing mechanism is linked to a hyperpolarization of the TMV (−66 mV), which leads to a block of the If and ICaL channels which ceases the automaticity of the SAN cell at the SEP entry. Fabbri et al. showed that the L-type calcium channel is one of the current responsible for the depolarization of the SAN (46). The blockage of the ICaL channel has been related to a slow down of the pacemaking function as described in different studies (38, 47).

Looking at the SAN-SEP structure, our model considers several characteristics that previous simulation studies did not include (7, 24, 25, 29). The heterogeneous electrophysiological characteristics of the SAN-SEP structure are represented as a combined gradient and mosaic model suggesting a necessary condition to elicit an AP in the SAN (pace) and have the excitation captured by the surrounding atrial myocardium (drive). Wilhelms et al. showed that the gradient model described the SAN heterogeneity in a 2D strand model better than the mosaic model and concluded that both gradient and mosaic models could not explain the increase of CV in the SAN periphery. They underlined the relevance of a 3D SAN model investigation (29). Similarly, Zhang et al. compared gradient and mosaic models in a 2D setting and concluded that the mosaic model was not able to reproduce the SAN behavior contrary to the gradient model (30). We showed that a combined gradient and mosaic model reproduced the SAN pace-and-drive behavior, with a rising CV in the SEPs (Figure 4C) composed of a mix of pacemaker and CT cells. CT myocytes were only present in the SEPs and not the SAN itself because the latter suppressed automaticity. This could explain why the mosaic models implemented in the 2D SAN models by Wilhelms et al. and Zhang et al., who for instance applied 41% of atrial cells in the SAN center based on the work of Verheijck et al. (48), were not able to reproduce the SAN behavior. Verheijck et al. also reported a share of 63% atrial cells in the SAN periphery but Dobrzynski et al. found a lower share of 26% (7). If we consider the junction of the nodal tissue and the CT (section 13 and and 14) as the periphery, our model includes a proportion of CT myocytes between 15% and 32%, which is consistent with the reported data. The percentage of CT myocytes then increases drastically in the SEP extension (75% to 95% within the last 0.15 mm of the SEP extension), making it challenging to delimit the periphery of the SAN in terms of CT cell percentage. This could also explain why a value of 63% of atrial cells in SAN periphery is given in literature (7, 30, 48), considering the increase of atrial cells from 75% to 95% within the three last 0.05 mm thick SEP extension sections.

The proposed model reproduced the experimentally reported CV increase in the SEPs ranging from 3 cm/s to 12 cm/s (2, 6, 24). Our model showed sinoatrial CV outside the LPS area that ranged between 9 cm/s to 5 cm/s before the SEP entry, where CV decreased to 1 cm/s due to the source-sink effect. However, the gradual CV increase in the transitional areas between SAN and the CT has not been clearly characterized experimentally, yet. In the 3D SEP model, we could quantify the CV increase from 20 cm/s at the SEP entry to >100 cm/s in the CT. In addition, we did not observed a complete suppression of SAN pacemaking in contrast to Fabbri et al. (21) who observed that the SAN was not able to pace due to the strong sink effect of the CT in a 1D strand with high conductivity.

Our model also included a realistic fiber direction, describing the gradual orientation change of the pacemaker cells towards the running CT as reported by Boyett et al. (6) and Dobrzynski et al. (7). Even if this assumed fiber direction could be refined and further studied, we clearly showed that this tissue property has a direct influence on the SEPs that preferentially activated the CT. This complements the observations by Fedorov et al. (2), who described atrial excitation of four coronary-perfused hearts via different sites corresponding to superior, lateral or inferior SEPs.

We proposed a modification of the mathematical model of the human atrial electrophysiology by Courtemanche et al. to reproduce the reported depolarization upstroke velocity of the CT (34). By increasing the conductance of the sodium channel as previously done in several studies (15, 29, 33) to reproduce experimentally measures repolarization times, we obtained a dV/dt_max_ of 319.49 V/s, which is 62% faster than in the rest of the RA. Future work could review the Courtemanche et al. (36) INa channel dynamics more in depth. The inactivation gates of the model might be re-parameterized following the methodology proposed for a ventricular model by Dutta et al. (49).

### Sinoatrial exit pathways: effects on activation

The effect of the width of SEPs was investigated in a 2D SAN model by Zyanterekov et al., who showed that narrower SEPs allowed for a more robust propagation to the surrounding atrial tissue (25). Our study confirmed this and provided further explanation by showing a decrease of the current propagating in the SEPs when their width was reduced (Figure 6C). Wider SEPs led to a higher influence of the CT on the SAN. We hypothesize that the gradient and mosaic distributions were no longer adjusted to maintain the drive capabilities under conditions of reduced current delivered by the wider SEP structures. Our results also suggest that a minimal width of the SEPs is required. The 3D SEP model provides as well a window to understand the influence of the SEP length (Figure 6C). Csepe et al. (3) described 2 mm to 3 mm long sinoatrial conduction pathways, connecting the SAN and the surrounding atrial tissue. In our model, SEPs shorter than 2 mm led to activation failure in the CT. We explain this by the fact that there were not enough SAN cells in the SEP to provide a smooth transition of the electrophysiological properties that would allow for capture of the excitation by the CT. Indeed, by increasing the SEP length, higher currents compared to the reference model propagated through the SEP and a CT activation was observed. These observations highlight the importance of the SEP dimensions, which modulate the hyperpolarizing effect of the CT cells on the SAN cells.

The CT activation occurred only via a single SEP in all simulations due to an exit block at the other SEPs (Figure 3B). This observation is in line with in vivo reports by Pambrun et al. who observed CT activation via only a single SEP in 49/50 human patients (13). The propagation was not possible via SEP n°4 and n°5 due to the default fiber direction implemented. Fiber direction rotated progressively from the SAN center towards the CT counterclockwise, favoring the propagation direction towards SEP n°1, n°2 and n°3 due to higher effective conductivity. CT activation happened most often via SEP n°1, n°2 or n°3, as described in vitro by Fedorov et al. and in vivo by Kharbanda et al., who reported atrial excitation via different exit locations (2, 50). We noticed that depending on the activation time of the SAN, different SEPs were activated. A faster SAN activation (160.80±3.42 ms) was related to the lateral SEP activation (SEP n°3) while longer SAN activation (211.20±27.05 ms) corresponded to activation of inferior SEPs (n°1 and n°2). Consistently with the observation of Fedorov et al., who described an atrial activation through inferior SEPs after a longer SAN conduction time than for superior or lateral SEP atrial activation, we provide an additional explanation to the location of atrial breakthrough sites (2).

### Leading pacemaker site

Robust SAN pace-and-drive capacity depends on the ability to shift the LPS to avoid exit block and overcome the source-sink mismatch (2, 6, 7, 11, 18, 23, 24, 39, 51), combined with the variability of the default LPS location from one patient to another (52). However, the reasons that lead to a LPS shift remained largely unclear and Kharche et al., for instance, modeled this phenomenon by displacing the LPS arbitrarily to different locations (24). Our model reproduced this phenomenon as a consequence of the natural rearrangement of the source-sink match, bringing a first explanation to the LPS location. The LPS was indeed situated in areas more remote from the SEPs, thus in areas where the CT cells had a lower influence on the SAN cells. Moreover, the LPS was never located in the SAN extremities, likely because of the too small available volume due to the ellipsoidal SAN shape. This could explain why the SAN head, the bigger part of the SAN structure located opposite of the CT and the SEPs was often observed as the LPS (9, 28, 53).

### Limitations

Fibrotic and adipose tissue surrounding the SAN was modeled by a cavity in the monodomain setting. However, the monodomain model has been proven to be sufficient to study depolarization phenomena of cardiac tissue (54). Kharche et al. provided a more realistic insulating border of the SAN, modeled by a thin connective tissue layer, and investigated the effect of fibrosis in the SAN (24). However, our approach of modeling a non-conducting cavity that insulates the SAN-SEP from the hyperpolarizing effect of the CT is sufficient to represent the effect of fibrosis as modeled by Vigmond et al. (55). Furthermore, to understand the effect of the insulating border, a bidomain model can be used, which will allow studying the effect of the thickness of the insulating border and its composition. Furthermore, our model could be further refined as proposed by Oren et al. (27) including different cellular models, for example, a mixture of myofibroblast and collagen (56, 57) to observe the cellular heterogeneity effect on the pace-and-drive mechanism of the SAN-SEP-CT structure.

The fiber direction implemented in our model was based on an interpretation of the SAN cell orientation from different studies (6, 7) rather than being directly derived from SAN tissue samples. Similarly, the microstructure of the exit pathways and transitional areas between the SAN and the surrounding cardiac tissue has not been definitely characterized experimentally. Our model considers these two aspects in an idealized geometrical model based on the sparse data reported from different studies (3, 8, 23). Even if our model allows defining the location of each SEP on the SAN surface, 5 locations were fixed in this study. The presence of single SEPs affected the LPS location, which could be further investigated in a 3D anatomical atrial model with varying SEP locations.

## CONCLUSION

In silico experiments underlined the importance of the SEP geometry, number and location for the SAN pace-and-drive mechanism to excite the atrial tissue. A combined gradient and mosaic model of the electrophysiological heterogeneity of the SAN-SEP-CT structure allows to overcome the source-sink mismatch. Additionally, changing the number and location of SEPs allowed for the first time to observe in silico a shift in the location of the LPS away from the hyperpolarized tissue to the best of our knowledge. The human SAN-SEP-CT model was able to pace and drive the RA at a physiological CL. These insights and the novel tool will inform future wet lab studies and provide the means for extensive in silico characterization of the heart’s natural pacemaker in health and disease.

## AUTHOR CONTRIBUTIONS

A.A., J.S., A.L. conceived the presented work and designed the research. A.A., J.S. carried out simulations, analyzed the data. A.A. and J.S. interpreted the results and drafted the manuscript. All authors provided critical feedback, approved and contributed to the final manuscript.

## ACKNOWLEDGMENTS

This work was supported by the Deutsche Forschungsgemeinschaft (DFG, German Research Foundation) – Project-ID 258734477 – SFB 1173 and the European High-Performance Computing Joint Undertaking EuroHPC under grant agreement No 955495 (MICROCARD) co-funded by the Horizon 2020 programme of the European Union (EU), the French National Research Agency ANR, the German Federal Ministry of Education and Research, the Italian ministry of economic development, the Swiss State Secretariat for Education, Research and Innovation, the Austrian Research Promotion Agency FFG and the Research Council of Norway. We gratefully acknowledge support by the state of Baden-Württemberg through bwHPC.

## Notes

### Competing Interest Statement

The authors have declared no competing interest.

